# Structural basis for intrinsic transcription termination

**DOI:** 10.1101/2022.08.30.505822

**Authors:** Linlin You, Expery Omollo, Chengzhi Yu, Rachel A. Mooney, Jing Shi, Liqiang Shen, Xiaoxian Wu, Aijia Wen, Dingwei He, Yuan Zeng, Yu Feng, Robert Landick, Yu Zhang

## Abstract

Efficient and accurate termination is required for gene transcription in all living organisms. Cellular RNA polymerases (RNAP) in both bacteria and eukaryotes can terminate their transcription through a factor-independent termination pathway (called intrinsic termination transcription in bacteria), in which RNAP recognizes terminator sequences, stops nucleotide addition, and releases nascent RNA spontaneously. Here we report a set of single-particle cryo-EM structures of *E. coli* transcription intrinsic termination complexes representing key intermediate states of the event. The structures show how RNAP pauses at terminator sequences, how the terminator RNA hairpin folds inside RNAP, and how RNAP rewinds the transcription bubble to release RNA and then DNA. These macromolecular snapshots define a structural mechanism for bacterial intrinsic termination and a pathway for RNA release and DNA collapse relevant for factor-independent termination by all RNA polymerases.

---

Genomic DNA is composed of functional genes transcribed to RNAs with defined 5′ and 3′ boundaries. The 5′ boundary is defined by a promoter, from which RNAP initiates RNA synthesis^1^. The 3′ boundary is defined by a terminator, at which RNAP stops transcription and releases the nascent RNA^2-5^. Efficient and accurate termination is required for gene transcription in all living organisms to ensure precise control of transcription units and to define the 3′ boundary of RNA transcripts^3,4^. Programmed transcription termination of RNAP occurs by two pathways, factor-independent (or intrinsic) and factor-dependent termination. Intrinsic termination requires only interactions of RNA and DNA with RNAP and can be modulated by transcription factors^5-8^. Intrinsic termination is prevalent in viruses and bacteria and used in eukaryotes by RNAPIII^6,8,9^.

The intrinsic terminator sequences in bacteria comprise a G/C-rich inverted repeat immediately followed by a stretch of thymidine that encodes an RNA with a G/C-rich terminator hairpin immediately followed by a uridine tract^10,11^. Genetic, biochemical, and single-molecule studies all suggest that intrinsic termination occurs via three sequential intermediates: (*i*) a pause state when RNAP pauses at the terminator; (*ii*) a hairpin-nucleation state when the RNA hairpin partially folds into RNAP; and (*iii*) a hairpin-completion state when the RNA hairpin completes folding and releases nascent RNA aided by the weak rU–dA base pairs. However, how the compact terminator sequence pauses, destabilizes, and dissociates RNAP remains elusive. To address the questions, we determined a set of cryo-EM structures representing the key intermediate states formed during intrinsic termination.

## RNAP pauses at the terminator sequence by adopting a half-swiveled conformation

RNAP first pauses at the –2/–1 positions (–1 corresponds to the position of the RNA 3′ end) of intrinsic terminators to begin intrinsic termination^8,12^. To study the structural mechanism of the terminator-induced pause, we reconstituted a paused transcription termination complex (TTC-pause) using *E. coli* RNAP core enzyme and a nucleic-acid scaffold that comprises a nascent RNA with λt_R2_ terminator sequence but lacking the upstream half of the hairpin stem (Fig. 1A). The TTC-pause structure was determined at 3.6-Å resolution using single-particle cryo-EM (Fig. S1 and Table S2). The cryo-EM map of TTC-pause shows unambiguous signals for ribonucleotides and deoxyribonucleotides within and near the transcription bubble, including the 10-bp RNA–DNA hybrid, the 10-nt single-stranded non-template DNA (ntDNA), the 12-bp downstream double-stranded (dsDNA), and 3-bp of the upstream dsDNA (Figs. 1B and 1C, and Movie S1).

**Figure 1.**
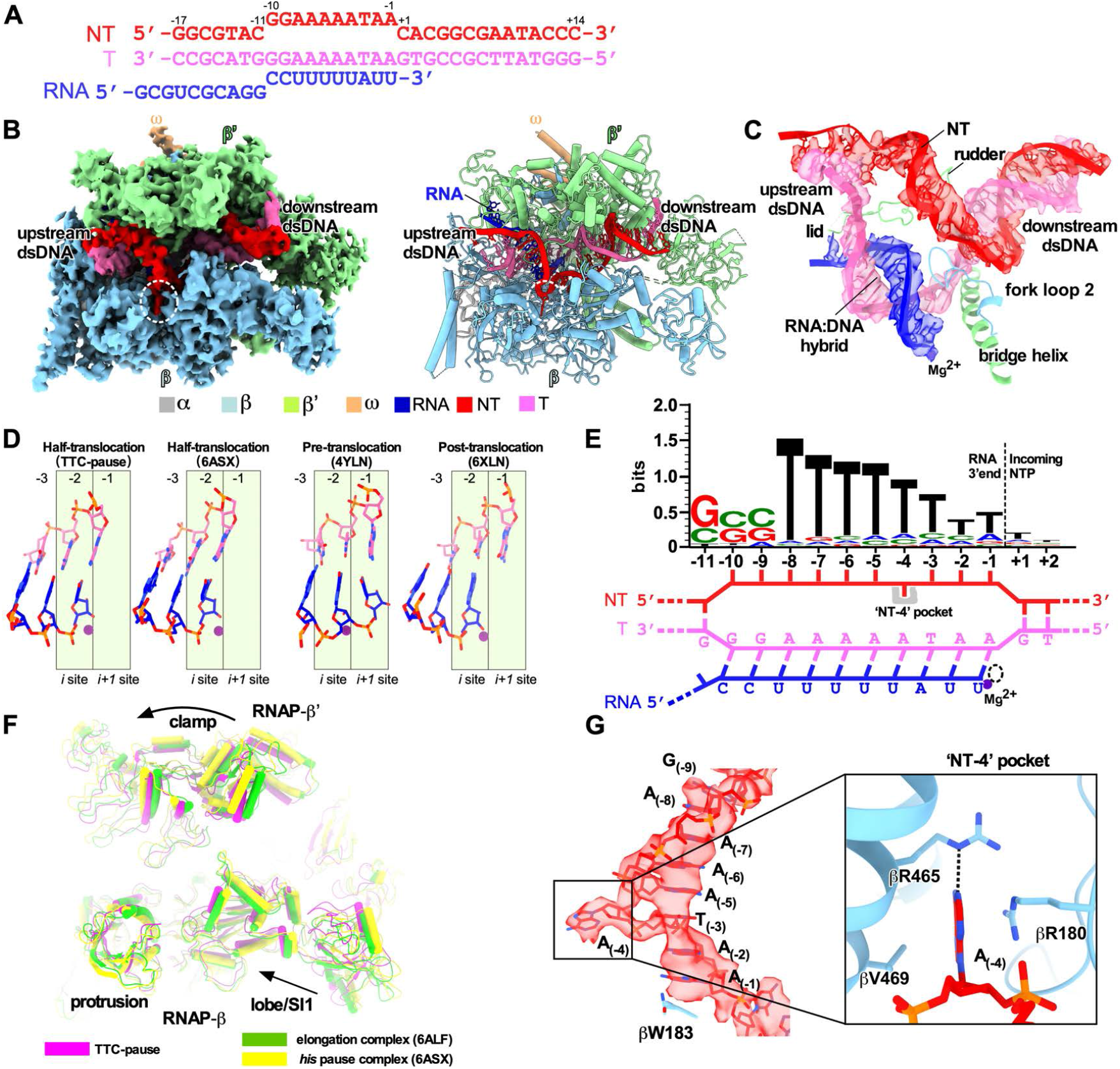
The TTC-pause complex. **(A)** The sequences of the nucleic-acid scaffold used for cryo-EM study. **(B)** The cryo-EM map (left) and structural model (right) for TTC-pause complex. The RNAP subunits and nucleic-acid chains are colored as in the color scheme. **(C)** The cryo-EM map for the nucleic-acid scaffold. **(D)** The comparison of the RNA–DNA translocation states among TTC-pause complex, half-translocation state in a hairpin-stabilized paused transcription elongation complex (*his*PEC; PDB: 6ASX), a pre-translocation complex (PDB: 4YLN), and a post-translocation complex (PDB: 6XLN). **(E)** The consensus sequence of bacterial intrinsic terminators (upper panel)^11^ and the schematic presentation of the nucleotides of the transcription bubble in the TTC-pause complex. **(F)** The conformational comparison among TTC-pause complex, TEC (PDB: 6ALF), and *his*PEC (PDB: 6ASX). **(G)** The cryo-EM map and structural model of ntDNA of the transcription bubble. The insert shows the interaction between the flipped A_(–4)_ nucleotide and the ‘NT–4’ pocket.

The RNA–DNA hybrid of TTC-pause adopts a half-translocated state, an off-pathway state that has been reported previously in paused transcription complexes^13-15^. In the half-translocated state, the RNA strand has translocated by 1-nt step from the pre-translocation state but the next template strand DNA (tDNA) nucleotide remains base-paired in the downstream dsDNA (Figs. 1D and S2C, and Movie S1). This bp is unwound and untranslocated except for slight positional shift of backbone phosphates (Fig. S2C). Half-translocation results in a tilted RNA–DNA hybrid and an empty NTP-binding site (i+1 site) lacking the template nucleotide for base-pairing to the incoming NTP (Fig. 1D). A survey of 100 native intrinsic terminators shows that intrinsic terminators resemble the consensus sequence for elemental pausing at two out of the three key positions (S_-10_U_-1_ *vs*. S_-10_Y_-1_G_+1_; Figs. 1E, S2A, and S2B)^11,16,17^. We infer that the sequences at the upstream and downstream edges of the transcription bubble (S_-10_, U_-1_) likely account for formation of the half-translocated conformation of TTC-pause complex^5^.

We also observed a global conformational change of RNAP involving multiple structural modules in TTC-pause compared with the normal transcription elongation complex (EC)^18^. The clamp module of RNAP rotates along an axis parallel to the bridge helix. The rotation movement of clamp module resembles the ‘swivel’ movement observed in the hairpin-stabilized paused elongation complex (PEC)^13,14,19^, but it only reaches halfway to the fully swiveled position (Fig. 1F). However, different from the swivel movement in the hairpin-stabilized PEC complex, the RNAP-β lobe/SI1 module in TTC-pause moves slightly towards the RNAP main cleft, likely due to unique interactions between RNAP and ntDNA of the transcription bubble (Fig. 1G and Movie S1). These movements of structural modules in TTC-pause likely accommodate the slightly shifted downstream dsDNA in the downstream dsDNA channel and stabilize the half-translocated RNA–DNA hybrid in the main cleft (Figs. S2C and S2D).

Our cryo-EM map of TTC-pause revealed a novel path of the single-stranded ntDNA in the transcription bubble across the RNAP main cleft. The ntDNA was accommodated in the upper floor of the main cleft and was separated from the active-site tunnel by the RNAP-β′ rudder loop and fork loop 2 (Figs. 1C and S2E). Eight nucleotides of the ntDNA form a continuous base stack that starts upstream at the RNAP-β′ coiled-coil (Fig. 1G and S2E), travels across the cleft to the RNAP-β protrusion and lobe, and follows fork loop 2 to reach the downstream dsDNA (Fig. S2E). The –4A is flipped out ∼180° from this continuous base stack. The backbone phosphates of the stacked nucleotides are stabilized by polar contacts with RNAP residues β′R271, β′R297, βR470, and βR473 (Fig. S2F). Residue βW183 supports the 8-nt base stack and separates it from the downstream dsDNA (Fig. S2F).

The interaction between RNAP and the –4 nucleotide is notable. The flipped out –4 nucleotide inserts into a protein pocket (NT–4 pocket) located in the gap between RNAP β protrusion and lobe (Fig. 1G). The A–4 base was sandwiched between residues βR180 and βV469; and the N1 atom of the adenine moiety makes a H-bond with βR465 (Figs. 1G and S2F). To test for potential effects of the pocket interaction on intrinsic termination, we measured the termination efficiency of RNAP derivatives bearing substitutions in the pocket at three different terminators. The results revealed no obvious effects on termination efficiency of disruption of either the stacking interaction (βR180A/βV469A) or H-bond interaction (βR465A) (Figs. S3A-F). Disruption of the pocket interaction also had no effect on transcriptional pausing at a hairpin-less λP_R_ terminator (Fig. S3G). Although no significant role in intrinsic termination was found, the pocket interaction also has been observed in a promoter escape complex and thus could play roles in other stages of transcription^20^.

## RNA hairpin nucleation in the RNA exit channel prepares the TTC for transcription bubble rewinding

The RNA hairpin of intrinsic terminators starts to fold in the RNA exit channel after RNAP pauses at terminators, a stage named hairpin nucleation^12,21^. To trap the hairpin-nucleation complex of transcription termination (TTC-hairpin), we incubated the TTC-pause complex with an antisense RNA10 (asRNA10) to form a 10-bp RNA duplex with the nascent RNA that mimics the hairpin stem (−20 to -11) and determined a TTC-hairpin structure at 3.1 Å resolution (Figs. 2A and S4, and Table S2). The cryo-EM map of TTC-hairpin revealed clear and sharp signals for the transcription bubble. The RNA–DNA hybrid adopts the same half-translocated conformation as in TTC-pause and the ntDNA nucleotides stack together and make interactions with RNAP essentially the same as in TTC-pause (Figs. 2C and S5A). The cryo-EM map of TTC-hairpin shows unambiguous signal for the 14-bp downstream dsDNA, the 3-bp upstream dsDNA, and, most importantly, the 10-bp dsRNA in the RNA exit channel (Figs. 2B and 2C, and Movie S2). These features indicate we successfully trapped the TTC-hairpin complex of intrinsic termination.

**Figure 2.**
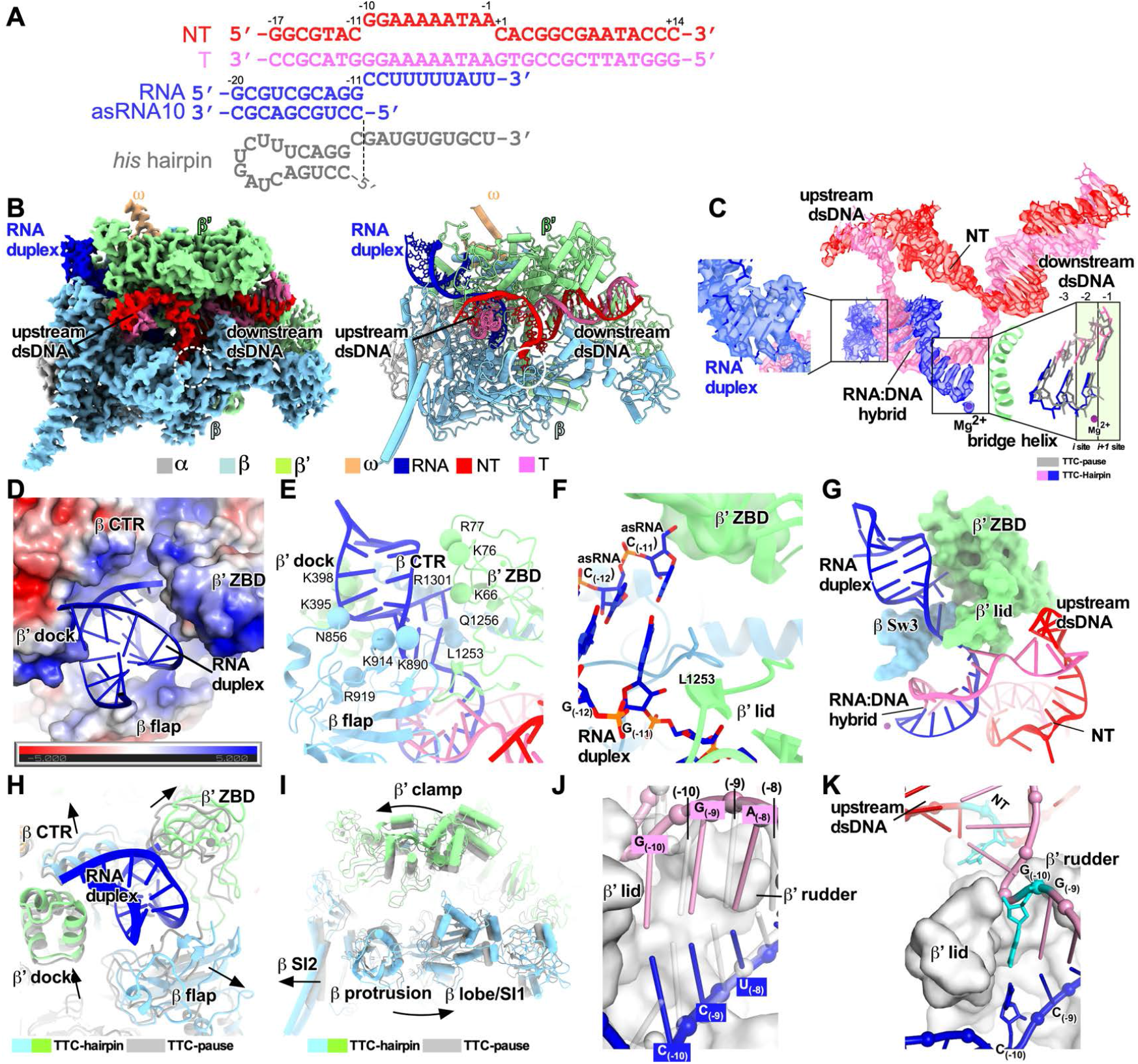
The TTC-hairpin complex. **(A)** The nucleic-acid scaffold and the antisense RNA (asRNA) used for cryo-EM structure determination. **(B)** The cryo-EM map (left) and structural model (right) for TTC-hairpin complex. **(C)** The cryo-EM map for nucleic-acid scaffold. The left insert shows the map for the RNA duplex in the RNA exit channel and the right insert shows the superimposition of the base pairs (–3 to 1) of the RNA–DNA hybrid between TTC-hairpin (colored as in the scheme) and TTC-pause (gray). **(D)** RNA duplex in the RNA exit channel. The electrostatic potential surface of RNAP was generated using APBS tools in Pymol. **(E)** The detailed interaction of the RNA duplex with residues in the RNA exit channel. Spheres, the Cβ atom of polar residues in H-bond distance with the phosphate backbone of RNA duplex. **(F)** The interaction of the –11 base pair of RNA duplex with residues in the RNA exit channel. **(G)** Further extension of RNA duplex is blocked by RNAP β′ ZBD, β′ lid, and β Sw3 motifs. **(H)** The structural comparison of RNA exit channel between TTC-pause (gray) and TTC-hairpin (colored). **(I)** The global conformational movement of TTC-hairpin (colored) compared with TTC-pause (gray). **(J)** The comparison of the first three base pairs of the RNA–DNA hybrid between TTC-hairpin (pink and blue) and TEC (gray) (PDB: 6ALF). The two structures were superimposed based in the RNAP-β′ lid and rudder motifs. **(K)** The G_(–10)_ of the tDNA is under the tunnel formed by RNAP-β′ lid and rudder and ready to pair with the –10 nucleotide of ntDNA.

The 10-bp dsRNA was tightly accommodated in the RNA exit channel by RNAP β′ dock, β′ C-terminal helix (β′ CTH), β′ zinc-binding domain (β′ ZBD), and β flap (Fig. 2D), consistent with the important roles of these elements in intrinsic termination^22,23^. The phosphate backbones of the RNA hairpin were held by RNAP polar residues in the RNA exit channel, including β′K66, β′K76, β′R77, β′Q1256, β′K395, β′K398, βN856, βK890, βK914, and βR919 (Fig. 2E). The innermost base pair of the RNA duplex reaches the bottom of the RNA exit channel. The –11G of the nascent RNA makes a base-stack interaction with βL1253 and the sugar moiety of the asRNA –11C fits into a shallow pocket of β′ ZBD (Fig. 2F). A positively charged groove likely guides ssRNA upstream of hairpin stem out of the RNA exit channel (Fig. S5B). Further extension of the RNA hairpin stem was stopped by β Sw3, β′ ZBD, and β′ lid (Fig. 2G). Most of these features of the hairpin stem closely match those observed for the hairpin-stabilized *his*PEC even though the TTC-hairpin stem is extended 1 bp closer to the RNA–DNA hybrid^13,14^.

The formation of an RNA duplex in TTC-hairpin causes a global conformational change of RNAP. The RNA duplex enlarges the RNA exit channel by stretching the four surrounding structural elements (Fig. 2H), likely triggering a global conformational change of RNAP structure modules, including further swiveling of the clamp module and outward rotation of the protrusion/lobe/SI1 module (Fig. 2I and Movie S2). The net result of such global rearrangement widens the main cleft and secures the hairpin in the RNA exit channel. Moreover, the fully swiveled conformation prevents trigger loop from refolding and further stabilizes the half-translocated RNA-DNA hybrid^13,24^. In short, our TTC-hairpin structure show that the RNA duplex in the RNA exit channel induces further conformational changes of RNAP compared with TTC-pause to stabilize the pause and likely to prepare for subsequent completion of RNA hairpin.

The completion of RNA hairpin requires melting of at least two upstream base pairs of RNA-DNA hybrid ^25^. Our cryo-EM structure of TTC-hairpin provides a structural clue for how the first base pair of the RNA–DNA hybrid is disrupted. In the TTC-hairpin structure, the half-translocated RNA–DNA hybrid shifts each of template nucleotide towards upstream from their respective positions in the pre-translocation state (Fig. 2J). As a result, the –10 nucleotide of tDNA shifts ∼1-bp register out of the RNAP active-center cleft to a new position under the small tunnel formed by the rudder and lid loops (Figs. 2K, S5C, and S5D). This new location allows unpairing of tDNA with the –10 nucleotide of the nascent RNA and its flipping through the tunnel to pair with the –10 nucleotide of ntDNA (Fig. S5E and Movie S3). Rewinding of this upstream base pair of the transcription bubble would weaken the next base-pair in the RNA–DNA hybrid due to loss of stacking interactions, thus enabling its disruption and lowering the energy barrier to RNA hairpin completion. The base pair at the –10 position shows the weakest map signals among the nucleotides of the RNA–DNA hybrid (Fig. S5F), indicative of its high flexibility necessary for spontaneous unwinding.

## Rewinding of transcription bubble promotes RNA release

The last stage of transcription termination involves disruption of RNA–DNA hybrid, completion of RNA hairpin, rewinding of transcription bubble, and release of RNA and DNA^8,12,25-28^. To trap intermediates of the last stage of intrinsic termination, we took advantage of asRNA-induced termination (Fig. 3A)^8^. The in vitro transcription termination assay suggests that, upon asRNA challenge, RNAP began to release nascent RNA in 30 seconds and released most of the nascent RNA at 5 minutes (Fig. 3B). Therefore, we challenged the TTC-pause complex with excess amount of 12-nt asRNA and vitrified the reaction mixture after 3-minute incubation, followed by cryo-EM data collection. A cryo-EM map was reconstructed at 3.5 Å from 54,471 particles (19% of total particles) after two rounds of 3D classification (Fig. S6 and Table S2). The cryo-EM density map for this novel complex (TTC-release) shows clear signals for 14-bp dsDNA in the downstream DNA channel, 6-bp upstream dsDNA located between the β′ clamp helices and β protrusion, and a partially rewound transcription bubble evidenced by the 5-nt single-stranded ntDNA (Figs. 3C and 3D, and Movie S4). The cryo-EM map shows no signal in the RNA-exit channel and weak disconnected signal in the active-center cleft that is likely from the non-specific rebinding of released RNA (Fig. 3D)^29^. The map features suggest that we have trapped an intermediate state of DNA–RNA release (TTC-release), in which RNA is released from the RNA exit channel, the transcription bubble is disrupted, and the ntDNA and tDNA of the transition bubble are partially rewound (Fig. 3E and Movie S4).

**Figure 3.**
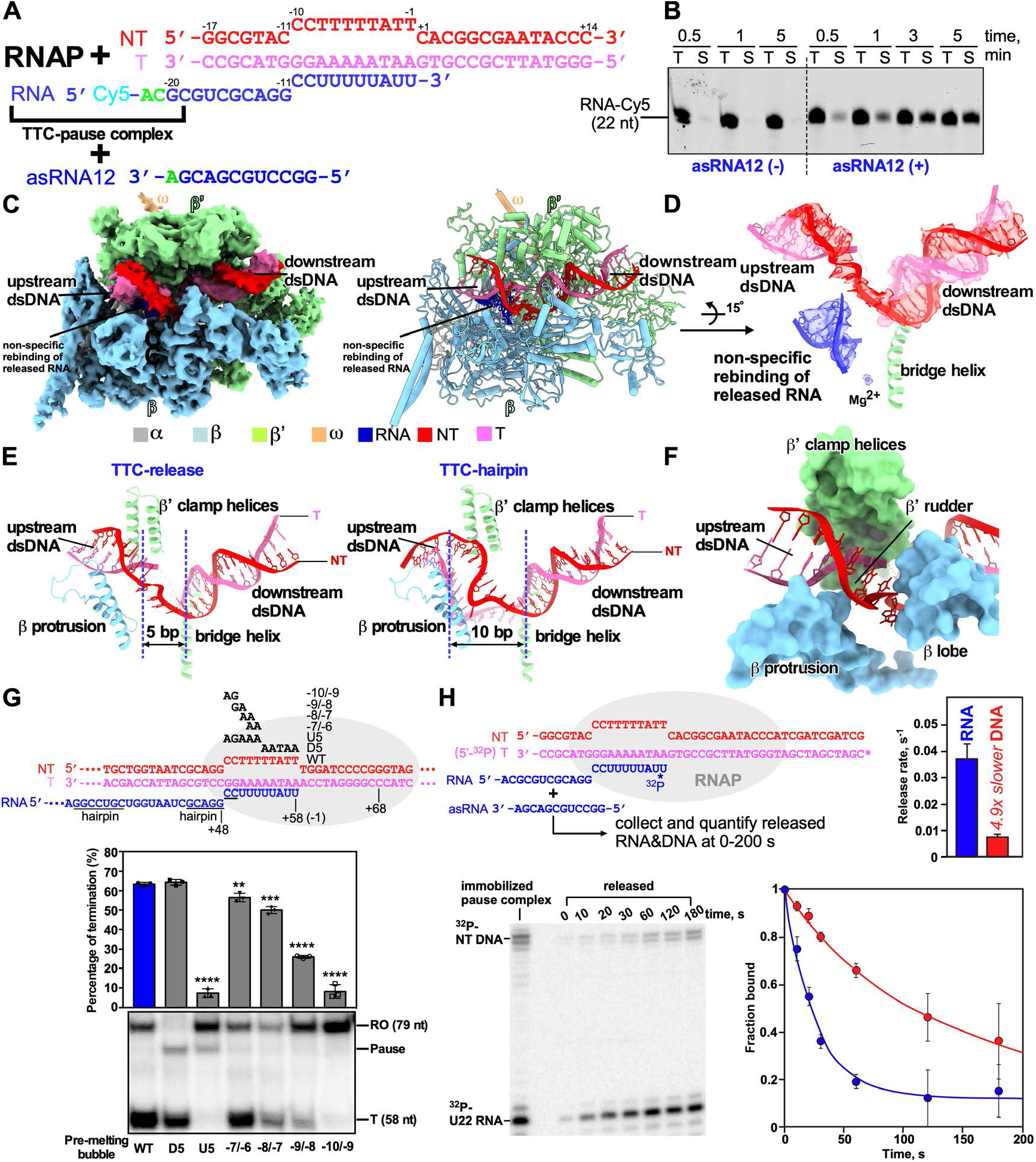
The release complex of transcription termination. **(A)** The asRNA-induced termination strategy used for obtaining the cryo-EM structure of TTC-release. **(B)** The results of RNA release assay show that asRNA12 induces release of nascent RNA in a time-dependent manner. T, total fraction; S, supernatant fraction. **(C)** The cryo-EM map and structural model of TTC-release. **(D)** The cryo-EM map and model of the nucleic-acid scaffold TTC-release. **(E)** The comparison of transcription bubble sizes of TTC-release and TTC-hairpin. **(F)** The rewound 5-bp dsDNA (–10 to –6) were loosely restrained by β protrusion, β′ clamp helices, and β′ rudder and further rewinding of the upstream dsDNA is stopped by the β lobe domain due to a closed clamp. **(G)** The *in vitro* transcription assay suggests that DNA unwinding is required for intrinsic termination at the λ_tR2_ terminator. Data are presented as mean ± SEM, n=3 biologically independent experiments. **P<0.01, ***P<0.001, ****P<0.0001. **(H)** RNA releases from TTC during termination faster than DNA releases. A TTC scaffold making all RNAP contacts with ^32^P label on the T strand 5′ O and RNA 3′ phosphodiester was immobilized on beads. Upon asRNA addition, the rate of RNA release was 0.037 ± 0.006 s^−1^ whereas DNA released at 0.0075 ±0.001 s^−1^.

The upstream dsDNA of TTC-release undergoes a radical conformation change compared with that of TTC-hairpin. The helix axis of the upstream dsDNA of TTC-release rotates ∼75° in an upward direction compared with that of TTC-hairpin (Fig. S7A). The rotation of the dsDNA relocates the 6-bp upstream dsDNA (–17 to –12) away from RNAP and thus becomes disordered in the structure (Fig. S7B). The rewound 5-bp dsDNA (–10 to –6) were loosely restrained by β protrusion, β′ CC, and β′ rudder (Fig. 3F). Further rewinding of the upstream dsDNA is stopped by the β lobe domain and requires cleft opening to a greater extent (Fig. 3F). The 5-nt single-stranded ntDNA remains in the same location as in TTC-pause and TTC-hairpin (Fig. 3E). The 14-bp dsDNA of TTC-release locates at the downstream dsDNA channel and makes interactions with RNAP as it does in an EC (Fig. S7C). Structural comparison shows that the RNAP clamp and lobe adopt the closed conformation seen in canonical EC (Fig. S7D), consistent with the previous findings that clamp opening is required for DNA release^30^. In short, our TTC-release structure suggests that DNA duplex remains bound to RNAP in a partially rewound form after RNA release.

The TTC-release complex also shows that transcription bubble rewinding is accompanied by hairpin completion and release of the nascent RNA^31,32^. The structure of the TTC-hairpin complex reveals the -10 nucleotide of tDNA is ready to unpair with RNA and to rewind with the -10 nucleotide of ntDNA (Figs. 2J, 2K and S5C-E). These results lead us to hypothesize that rewinding of the transcription bubble triggers the subsequent steps of hairpin completion and RNA release. To test our hypothesis, we measured termination efficiency at λ_tR2_ with pre-melted regions at various positions close to the termination site (Figs. 3G and S8A). The results show that pre-melting the downstream half of the transcription bubble (–5 to –1; D5) has no effect on the termination at all, but pre-melting the upstream half of the transcription bubble (–10 to –6; U5) completely abolished termination (Fig. 3G). This result supports the ideas that rewinding of transcription bubble is required for intrinsic termination and that the process initiates from the upstream end, consistent with a previous report^32^. We next performed a more refined mapping to define the minimal requirements of bubble rewinding for efficient termination. Pre-melting two base pairs in the middle of the transcription bubble (–7/–6) had little effect. Moving the 2-bp pre-melted region upstream gradually decreased the termination efficiency. Strikingly, pre-melting the first two base pairs (–10/–9) completely abolished termination, highlighting the crucial role of rewinding the first two base pairs (–10/–9) during intrinsic termination (Fig. 3G). The same results were obtained using the ϕt_500_ terminator sequence (Fig. S8B). These results support our hypothesis that rewinding the transcription bubble is an important step for intrinsic termination, likely through reducing the energetic barriers for hairpin completion and RNA release. We also noticed that termination occurred, albeit with low efficiency, in response to asRNAs (asRNA –11 and –10) that only form a duplex in the RNA exit channel and do not disrupt the RNA–DNA hybrid (Fig. S8C)^33^. This result suggests that rewinding the transcription bubble is sufficient to induce termination even without hairpin extension, consistent with bubble rewinding beginning the RNA release process. Together, our cryo-EM structures and in vitro transcription results suggest that rewinding the upstream two base pairs of transcription bubble is a requisite for subsequent RNA hairpin completion and RNA release.

An intriguing implication of our structures is that the nascent RNA dissociates before RNAP releases dsDNA, consistent with evidence that terminated RNAP can remain associated with and slide on DNA after termination^34,35^. We tested this idea by measuring the rates of RNA and DNA release from TTC in physiologically relevant solutes using a scaffold that provided all RNAP–DNA contacts (Fig. 3H). After addition of asRNA to trigger termination, RNA was released ∼5 times faster than DNA. This result indicates that the post-termination, the binary RNAP–DNA complex survives after RNA release. This binary complex may dissociate more slowly *in vivo* where sliding RNAP would not immediately encounter a DNA end.

## Discussion

On the basis of our cryo-EM structures and biochemical evidence, we propose a detailed model for bacterial intrinsic termination (Fig. 4 and S9): (*i*) RNAP pauses at the terminator site containing elemental pause-like sequence motif (S_–10_U_–1_), where it adopts the ‘half-swiveled’ conformation that accommodates a half-translocated RNA-DNA hybrid in the active-site cleft; (*ii*) initial folding of terminator hairpin enlarges the RNA exit channel, induces a ‘full-swivel’ RNAP conformation, and weakens the upstream RNA–DNA base-pair interactions; (*iii*) RNAP rewinds the two base pairs of the upstream transcription bubble and clears the energetic barrier for subsequent RNA hairpin completion; (*iv*) the terminator hairpin completes formation in the RNA exit channel by pulling the RNA out the exit channel either by ‘hybrid shearing’ (RNA moves but tDNA does not) or by pulling the RNA–DNA hybrid upstream by ‘forward translocation’(the downstream duplex melts without nucleotide addition to RNA) (Fig. S9)^26,27^, meanwhile, rewinding of the transcription bubble propagates along with the RNA translocation; and (*v*) RNAP releases the nascent RNA but retains the partially rewound DNA^34,35^. RNAP may either slide on genomic DNA or finally dissociate DNA spontaneously or aided by pro-termination factors^5,34-38^.

**Figure 4.**
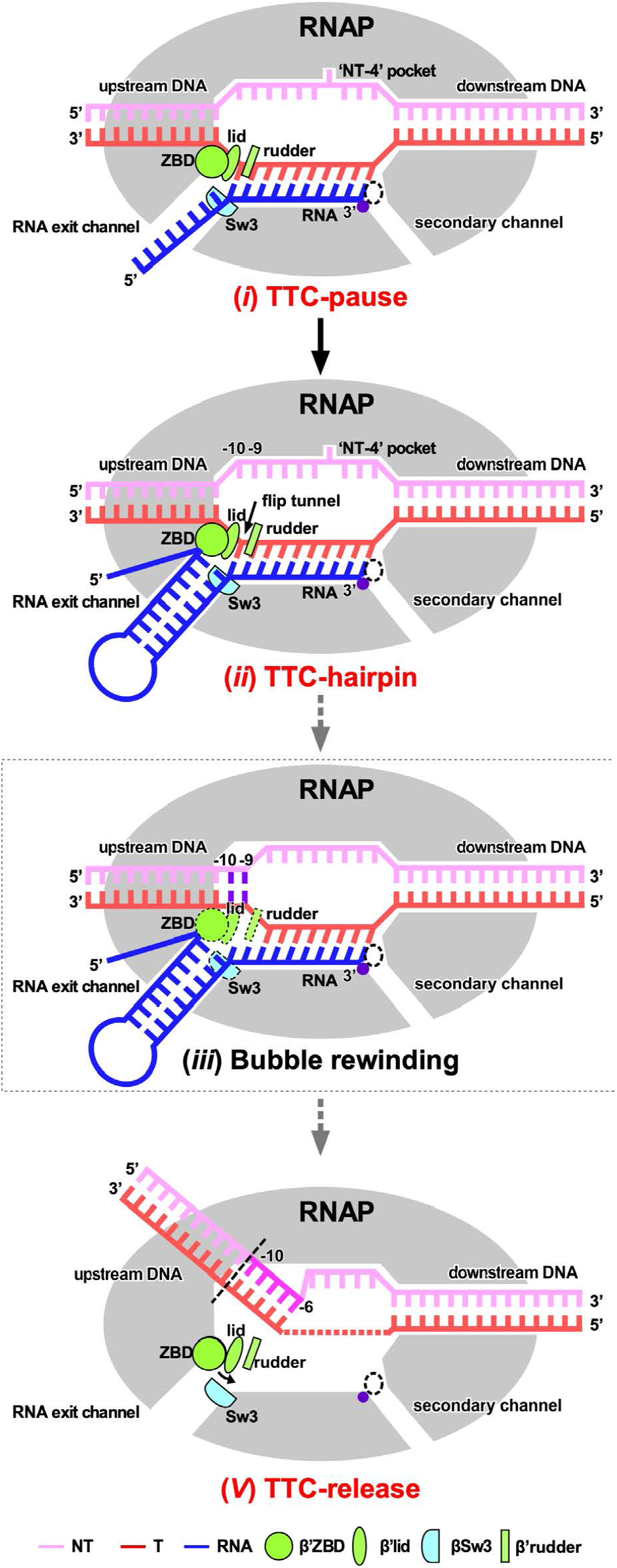
The proposed pathway of intrinsic termination. Intermediate states confirmed by structures obtained in this study are labeled in red text. A proposed intermediate state in the pathway is labeled in black text and boxed by dotted lines. See also Fig. S9.

In summary, our cryo-EM structures of transcription termination complexes reveal how bacterial RNAP pauses at intrinsic terminator sequence, how a terminator hairpin nucleates its folding in the RNA exit channel to weaken the RNA–DNA hybrid, how RNAP rewinds the transcription bubble to allow completion of the terminator hairpin and release of nascent RNA, and how RNAP retains the partially rewound dsDNA. The structures provide structural mechanisms for bacterial intrinsic termination. The “DNA rewinding-triggered RNA release” mechanism provides clues for factor-independent termination at hairpin-less terminator sequences by eukaryotic RNA polymerase III^39,40^.

## Methods

### Plasmid construction

Plasmid pET-28a-TEV-*Ec-*σ^70^ (Table S1) was constructed by inserting the *E. coli* σ^70^ gene amplified from *E. coli* genomic DNA into a modified pET-28a plasmid carrying the TEV protease cleavage site by a homogenous recombination method (pEASY®-Basic Seamless Cloning and Assembly Kit, Transgen Biotech, Inc.). Derivatives of pRL706 were constructed by primer-mediated, site-directed mutagenesis using NEB Q5 site-directed mutagenesis reagents.

### *E. coli* RNAP core enzyme

*E. coli* RNAP core enzyme for cryo-EM and most *in vitro* assays was over-expressed from *E. coli* BL21(DE3) (Novo protein, Inc.) carrying p*Ec*ABC and pCDF-*Ec* rpoZ (Table S1) and purified as described^41^. *E. coli* RNAP core enzyme used in experiments shown in Figs. 3H and S8C was purified as described previously^42^.

*E. coli* RNAP core enzyme with substitutions in the β subunit used in experiments shown in Fig. S3 was purified using β-subunit overexpression from pRL706 derivatives (Table S1). Each plasmid was independently transformed into *E. coli* strain RL1204 (Table S1) and a single colony was inoculated into 30 mL LB + 100 μg carbenicillin/mL and grown overnight at 37 °C. The saturated cell culture was added to 1 L fresh LB + 100 μg carbenicillin/mL and grown at 37 °C with adequate aeration by orbital shaking in a Fernbach flask until apparent OD_600_ reached 0.2. β subunit overexpression was induced by adding IPTG (Gold Biotechnology) to 1 mM and cell growth was monitored until apparent OD_600_ reached 0.9. The cells were harvested and homogenized by sonication in 30 mL Lysis Buffer (50 mM Tris-HCl, pH 7.9, 5% v/v glycerol, 233 mM NaCl, 2 mM EDTA, 10 mM β-mercaptoethanol, 100 μg PMSF/mL, 1 tablet of protease inhibitor cocktail (Roche), and 10 mM DTT). After removing cell debris by centrifugation (27000 × *g*, 15 min, 4 °C), DNA binding proteins including target RNAPs were precipitated by addition of polyethyleneimine (PEI, Sigma-Aldrich) to 0.6% (w/v) final and stirred for at least 10 min. After centrifugation (11000 × *g*, 15 min, 4 °C), the protein pellet was resuspended in 25 mL of PEI wash buffer (10 mM Tris-HCl, pH 7.9, 5% v/v glycerol, 0.1 mM EDTA, 5 μM ZnCl_2_, 500 mM NaCl) to remove non-target proteins. After centrifugation (11000 × *g*, 15 min, 4 °C), RNAP was eluted from the pellet into 25 mL PEI Elution Buffer (10 mM Tris-HCl, pH 7.9, 5% v/v glycerol, 0.1 mM EDTA, 5 μM ZnCl_2_, 1 M NaCl). The crude extract of RNAP was subjected to sequential FPLC purifications using Ni^2+^-affinity (HisTrap FF 5 mL, Cytiva), followed by purification using a Heparin column (HiTrap FF 5 mL, Cytiva). The purified RNAPs were dialyzed into RNAP Storage Buffer (10 mM Tris-HCl, 25% v/v glycerol, 100 mM NaCl, 100 μM EDTA, 1 mM MgCl2, 20 μM ZnCl2, 10 mM DTT). Samples were aliquoted, flash frozen in liquid nitrogen and stored in –80 °C.

### *E. coli* σ^70^

*E. coli* σ^70^ was over-expressed in *E. coli* BL21(DE3) cells (Novo protein, Inc.) carrying pET28a-TEV-*Ec-*σ^70^ (Table S1). Protein expression was induced with 0.3 mM IPTG at 18 ºC for 14 h when OD_600_ reached to 0.6-0.8. Cell pellet was lysed in lysis buffer B (50 mM Tris-HCl, pH 7.7, 500 mM NaCl, 5% (v/v) glycerol, 5 mM β-mercaptoethanol, and 0.1 mM PMSF) using an Avestin EmulsiFlex-C3 cell disrupter (Avestin, Inc.). The lysate was centrifuged (16,000 × *g*; 50 min; 4 ºC) and the supernatant was loaded on to a 2 mL column packed with Ni-NTA agarose beads (Smart-Lifesciences, Inc.). The bound proteins were washed by lysis buffer B containing 20 mM imidazole and eluted with the lysis buffer B containing 400 mM imidazole. The eluted fractions were supplemented with TEV protease and transferred to a dialysis bag to exchange buffer to 20 mM Tris-HCl, pH 7.7, 150 mM NaCl, 5% (v/v) glycerol, 5 mM β-mercaptoethanol. The sample was reloaded onto the Ni-NTA column, and the tag-free protein was retrieved from the flow-through fraction. The sample was diluted, loaded onto a Q HP column (HiPrep Q HP 16/10, Cytiva) and eluted with a salt gradient of buffer A (50 mM Tris-HCl, pH 7.7, 150 mM NaCl, 5% (v/v) glycerol, 1 mM DTT) and buffer B (50 mM Tris-HCl, pH 7.7, 500 mM NaCl, 5% (v/v) glycerol, 1 mM DTT). The fractions containing target proteins were collected, concentrated to 5.2 mg/mL, and stored at -80 ºC.

*E. coli* σ^70^ used in experiments shown in Figs. S3 and S8C was purified as described previously^42^.

### *E. coli* RNAP holoenzyme

*E. coli* RNAP core (3 μM, final concentration) and σ^70^ (12 μM, final concentration) were incubated in 0.5 mL 20 mM Tris-HCl, pH 7.7, 100 mM NaCl, 1% (v/v) glycerol, 1 mM DTT for 2 h at 4 °C. The reaction mixture was applied to a Superdex 200 10/300 column (Cytiva) equilibrated in 20 mM Tris-HCl, pH 7.7, 100 mM NaCl, 1% (v/v) glycerol, 1 mM DTT. Fractions containing RNAP holoenzyme were collected and concentrated to ∼5 mg/mL, and stored at -80 °C.

### Nucleic-acid scaffolds

Nucleic-acid scaffolds (Table S1) for cryo-EM study of *E. coli* TTC-pause and TTC-hairpin were prepared as follows: nontemplate-strand DNA (5’-GGCGTACGGAAAAATAACACGGCGAATACCC-3’; 0.3 mM, final concentration; Sangon Biotech), template-strand DNA (5’-GGGTATTCGCCGTGAATAAAAAGGGTACGCC-3’; 0.33 mM, final concentration; Sangon Biotech) and RNA (5’-GCGUCGCAGGCCUUUUUAUU-3’; 0.39 mM, final concentration; GenScript Biotech Corp.) in 50 μL annealing buffer (5 mM Tris-HCl, pH 8.0, 200 mM NaCl, and 10 mM MgCl_2_) were heated for 5 min at 95°C, cooled to 22 °C in 2 °C steps with 30 s per step using a thermal cycler.

Nucleic-acid scaffold (Table S1) for cryo-EM study of *E. coli* TTC-release was prepared as follows: template-strand DNA (5’-GGGTATTCGCCGTGAATAAAAAGGGTACGCC-3’; 0.39 mM, final concentration; Sangon Biotech) and RNA (5’-Cy5-ACGCGUCGCAGGCCUUUUUAUU-3’; 0.3 mM, final concentration; GenScript Biotech Corp.) in 50 μL annealing buffer (5 mM Tris-HCl, pH 8.0, 200 mM NaCl, and 10 mM MgCl_2_) were heated for 5 min at 95°C, cooled to 22 °C in 2 °C steps with 30 s per step using a thermal cycler.

For experiments shown in Figs 3G, S3, and S8C, DNA and RNA oligos (Table S1; Fig. S8C) were obtained from Integrated DNA Technologies (IDT; Coralville, IA) and were purified by denaturing polyacrylamide gel electrophoresis (PAGE; 15% 19:1 acrylamide: bisacrylamide, 45 mM Tris-borate, pH 8.3, 1.25 mM Na_2_EDTA, 8M urea) before use, unless otherwise stated. [γ-^32^P]ATP, [α-^32^P]UTP and [α-^32^P]GTP were obtained from PerkinElmer Life Sciences; rNTPs, from Promega (Madison, WI, USA); and 3’deoxy GTP (3′dGTP) from Jena Bioscience.

### Cryo-EM data collection and processing: *E. coli* TTC-pause

*E. coli* RNAP core enzyme (20 μM, final concentration) and the nucleic-acid scaffold (26 μM, final concentration) were incubated in 0.5 mL 10 mM HEPES, pH 7.5, 5 mM KCl, 5 mM MgCl_2_, 3 mM DTT at room temperature for 50 min. The mixture was applied to a Superdex 200 10/300 column (Cytiva) equilibrated in 10 mM HEPES, pH 7.5, 5 mM KCl, 5 mM MgCl_2_, 3 mM DTT. Fractions containing TTC-pause complex were collected and concentrated to 13 mg/mL.

The freshly purified *E. coli* TTC-pause at 13 mg/mL was incubated with 3-([3-cholamidopropyl] dimethylammonio)-2-hydroxy-1-propanesulfonate (CHAPSO, 8 mM, final concentration; Hampton Research Inc.) prior to grid preparation. The complex (3 μL) was subsequently applied on a glow-discharged C-flat CF-1.2/1.3 400 mesh holey carbon grids (Electron Microscopy Sciences), blotted with Vitrobot Mark IV (FEI), and plunge-frozen in liquid ethane with 95% chamber humidity at 10 °C.

The data were collected on a 300 keV Titan Krios (FEI) equipped with a K2 Summit direct electron detector (Gatan) at National Center for Protein Sciences Shanghai. A total of 1,796 images were recorded using the Serial EM^43^ in super-resolution counting mode with a pixel size of 1.0 Å, and a dose rate of 8 electrons/pixel/s. Movies were recorded at 200 ms/frame for 7.6 s (38 frames total) and defocus range was varied between -1.2 μm and -2.2 μm. Frames of individual movies were aligned using MotionCor2^44^, and contrast-transfer-function estimations were performed using CTFFIND4^45^. About 1,000 particles were picked and subjected to 2D classification in RELION 3.0^46^. The resulting distinct two-dimensional classes were served as templates for particle auto-picking and 203,216 particles were picked out. The resulting particles were subjected to 2D classification in RELION 3.0 by specifying 100 classes^46^. Poorly-populated classes were removed. We used a 50-Å low-pass-filtered map calculated from structure of *E. coli* RNAP core enzyme (PDB: 6ALF) as the starting reference model for 3D classification (N=3). The final concentration maps calculated from 132,648 particles were obtained through 3D auto-refinement, CTF-refinement, Bayesian polishing, and post-processing in RELION 3.0. Gold-standard Fourier-shell-correlation analysis indicated a mean map resolution of 3.58 Å (Fig. S1)

The model of RNAP core enzyme from the cryo-EM structure of *E. coli* TEC (PDB: 6ALF) was manually fit into the cryo-EM density map using Chimera^47^. Rigid body and real-space refinement was performed in Coot^48^ and Phenix^49^.

### Cryo-EM data collection and processing: *E. coli* TTC-hairpin

The freshly purified *E. coli* TTC-pause complex (13 mg/mL, 33 μM, final concentration) and antisense RNA (asRNA10, 330 μM, final concentration) were incubated in 30 μL 10 mM HEPES, pH 7.5, 5 mM KCl, 5 mM MgCl_2_, 3 mM DTT for 3 h at 4°C. The sample was vitrified by the same procedure as the TTC-pause complex.

The data were collected on a 300 keV Titan Krios (FEI) equipped with a K2 Summit direct electron detector (Gatan) at center of Electron Microscopy, Zhejiang University. A total of 3,888 images were recorded using the Serial EM^43^ in counting mode with a pixel size of 1.307 Å, and a dose rate of 9.9 electrons/pixel/s. Movies were recorded at 250 ms/frame for 10 s (40 frames total) and defocus range was varied between -1.8 μm and -2.6 μm. Frames of individual movies were aligned using MotionCor2^44^, and contrast-transfer-function estimations were performed using CTFFIND4^45^. About 1,548 particles were picked and subjected to 2D classification in RELION 3.0^46^. The resulting distinct two-dimensional classes were served as templates and a total of 1,548 particles for were picked out. The resulting particles were subjected to 2D classification in RELION 3.0 by specifying 100 classes ^46^. Poorly-populated classes were removed. We used a 50-Å low-pass-filtered map calculated from structure of *E. coli* RNAP core enzyme^18^ (PDB: 6ALF) as the starting reference model for 3D classification (N=4). Two same classes were combined and 360,313 particles were used for 3D auto-refinement. To resolve heterogeneity around RNA hairpin in RNA exit channel, a soft mask that excludes RNA duplex and nearby protein regions (β′ ZBD, β′ dock, β flap, β CTR) was generated in Chimera and RELION 3.0. The mask was used to make a subtracted particle stack in RELION 3.0. The subtracted particles were applied for masked 3D classification (N=6, without alignment), the best-resolved class containing obvious RNA hairpin (293,294 particles) was used for 3D auto-refinement, CTF-refinement and Bayesian polishing, Postprocessing. Gold-standard Fourier-shell-correlation analysis indicated a mean map resolution of 3.05 Å (Fig. S4). The structural model of RNAP core enzyme from the cryo-EM structure of *E. coli his*PEC (PDB: 6ASX) was manually fit into the cryo-EM density map using Chimera^47^. Rigid-body and real-space refinement was performed in Coot^48^ and Phenix^49^.

### Cryo-EM data collection and processing: *E. coli* TTC-release

*E. coli* RNAP core enzyme (20 μM, final concentration) and nucleic-acid scaffold comprising tDNA and RNA (30 μM, final concentration) were incubated in 0.5 mL 10 mM HEPES, pH 7.5, 50 mM KCl, 5 mM MgCl_2_, 3 mM DTT at room temperature for 30 min. The ntDNA (200 μM, final concentration) was subsequently added and the mixture was further incubated at room temperature for 30 min. The mixture was applied to a Superdex 200 10/300 column (Cytiva) equilibrated in 10 mM HEPES, pH 7.5, 50 mM KCl, 5 mM MgCl_2_, 3 mM DTT. Fractions containing TTC-pause complex were collected and concentrated to 17 mg/mL (∼43 μM), then the sample mixed with CHAPSO (Hampton Research, Inc.) to a final concentration 8 mM, 430 μM (final concentration) antisense RNA12 (asRNA12) was added and the reaction was incubated for 3 min prior to grid preparation. The complex (3 μL) was quickly applied on a glow-discharged UltraAuFoil R1.2/1.3 300 mesh holey Au grids (Quantifoil Micro Tools GmbH), blotted with Vitrobot Mark IV (FEI), and plunge-frozen in liquid ethane with 100% chamber humidity at 22 °C.

The data were collected on a 300 keV Titan Krios (FEI) equipped with a K3 Summit direct electron detector (Gatan) at National Center for Protein Sciences Shanghai. A total of 1,355 images were recorded using the EPU using super-resolution counting mode for 2.67 s exposures in 40 frames to give a total dose of 49.65 electrons per Å^2^ with defocus range of - 1.2 to -2.2 μm. Frames of individual movies were aligned using MotionCor2^44^, and contrast-transfer-function estimations were performed using CTFFIND4^45^. About 1,000 particles were picked and subjected to 2D classification in RELION 3.0^46^. The resulting distinct two-dimensional classes were served as templates for auto-picking and a total of 592,713 particles were picked out. The resulting particles were subjected to 2D classification in RELION 3.0 by specifying 100 classes^46^. Poorly-populated classes were removed. We used a 50-Å low-pass-filtered map calculated from TTC-hairpin map as the starting reference model for 3D classification (N=6). Two classes were combined and 282,423 the particle numbers used for 3D auto-refinement. To resolve heterogeneity about the DNA and RNA in the main cleft, a soft map that excludes the upstream dsDNA, the transcription bubble, the RNA-DNA hybrid, the downstream dsDNA, and the β’clamp domain nearby was generated in Chimera and RELION 3.0. The mask was used to make a subtracted particle stack in RELION 3.0. The subtracted particles were applied for masked 3D classification (N=6, without alignment), the 3D class of TTC-release (54,309 particles) were used for 3D auto-refinement, CTF-refinement and Bayesian polishing, Postprocessing. Gold-standard Fourier-shell-correlation analysis indicated a mean map resolution of 3.48 Å (map A in Fig. S6D). The structural model of RNAP core enzyme from the cryo-EM structure of *E. coli his*PEC (PDB: 6ASX) was manually fit into the cryo-EM map using Chimera^47^. Rigid body and real-space refinement was performed in Coot^48^ and Phenix^49^. The other two major classes were also processed under the similar procedure resulting in two maps at 3.40 Å (map B and C in Fig. S6D), both of which show little signal for the upstream dsDNA, the non-template ssDNA, and the ssRNA in the RNA exit channel, although clear signals for the downstream dsDNA and the half-translocated RNA–DNA hybrid. These features suggest that these two complexes were likely not properly assembled during complex reconstitution.

### Fluorescence-detected RNA release assay

To study asRNA-induced RNA release, *E. coli* RNAP core enzyme (200 nM, final concentration) and TTC-release nucleic-acid scaffold (800 nM, final concentration) comprising Cy5-labelled RNA and tDNA were incubated in 300 mL transcription buffer (50 mM Tris-HCl, pH 8.0, 50 mM KCl, 5 mM MgCl_2_, 5 mM β-mercaptoethanol at room temperature for 15 min. Then ntDNA (2 µM, final concentration) was subsequently added and the mixture was further incubated at room temperature for 15 min to form TTC-pause complex. The complex was immobilized on 150 mL Ni-NTA agarose beads (smart-lifesciences, Inc.) pre-washed with transcription buffer. The immobilized complex was washed with 300 mL transcription buffer for three times. The reaction mixture was aliquoted and each of the aliquot (70 μL) was supplemented with 7 mL asRNA12 (final concentration: 1 μM) to induce RNA release. For each of the aliquots, 20 mL reaction mixtures were taken out as reference of total amount of nascent RNA. The resulting reaction mixtures (50 μL) were separated into supernatant and pellet fractions by centrifugation at specified time points. Both the total and supernatant samples (20 mL) were mixed with 5 μL loading buffer (8 M urea, 20 mM EDTA, 0.025% xylene cyanol), 95 °C boiled for 5 min, and cooled down in ice for 5 min. RNA were separated by 20% urea-polyacrylamide slab gels (19:1 acrylamide/bisacrylamide) in 90 mM Tris-borate, pH 8.0 and 0.2 mM EDTA and analyzed by fluorescein scanning (Typhoon; GE Healthcare, Inc.).

### Release assay to measure RNA and DNA rates release during intrinsic termination

The TTC release scaffold (Figure 3A) was modified by addition of 12 downstream base pairs to eliminate any effect of RNAP–DNA-end contacts that could affect the DNA release rate (Figure 3H). ECs were reconstituted one nucleotide upstream from the U8 termination site by incubation of 1 µM RNA21, 200 nM T DNA that was 5′ ^32^P-labeled by treatment with [β-^32^P]ATP polynucleotide kinase, and 400 nM *E. coli* core RNAP in 100 µL EC buffer (10 mM Hepes, pH 8.0, 50 mM KGlutamate, 10 mM MgOAc, 0.1 mM EDTA, 5 μg acetylated BSA/mL and 1 mM DTT) for 5 min at 37 °C. Non-template DNA (2 µM) was added and the mixture was incubated for an additional 5 min at 37 °C. The final ratio of RNAP:RNA:T:NT was 2:5:1:10. The estimated concentration of the assembled transcription termination complex was 200 nM. Heparin (50 µg/mL) was added to the mixture to prevent core RNAP from rebinding to the scaffold after release. The RNA was extended to U8 by reaction with 2 µM [β-^32^P]UTP (136 Ci/mmol) yielding TTC with 5′ ^32^P-labeled T DNA and 3′ ^32^P-labeled RNA. The complex was immobilized on 20 µL of Ni-NTA agarose beads (Qiagen) with occasional pipetting. After 10 min incubation at room temperature, the immobilized complex was washed with EC buffer (five cycles of centrifugation and resuspension of the pelleted beads in 200 µL of fresh EC buffer). The washed bead-complex was resuspended in 200 µL EC buffer. One 5 µL portion was mixed with 5 µL stop buffer (8 M urea, 50 mM EDTA, 90 mM Tris-Borate buffer, pH 8.3, 0.02% bromophenol blue, 0.02% xylene cyanol) for total 0-timepoint sample. A second 5 µL portion was incubated with ATP, CTP, and 3′-dGTP (150 µM each), then combined with stop buffer as a check for TTC integrity. (Fig. 3H). The magnetic Ni-NTA beads were pulled to one side using a magnet, then 25 µL of the supernatant was passed through a nitrocellulose microfilter plate (384 wells) placed on top of a multi-well plate vacuum manifold (Operated at 20 Hg) with a receiver plate on the bottom (Pall Corporation). A portion (5 µL) of the filtrate was mixed with 5 µL stop buffer for the released 0-timepoint sample. To initiate the termination reaction and monitor RNA and DNA release, asRNA – 12mer (Fig. 3H) was added to a final concentration of 1 µM and supernatant portions (25 µL) were removed after 10, 20, 30, 60, 120 and 180 s, filtered, and combined with stop buffer as described for the 0-timepoint sample. Samples were then analyzed by denaturing PAGE (15% 19:1 acrylamide: bis-acrylamide, 45 mM Tris-borate, pH 8.3, 1.25 mM Na_2_EDTA, 8M urea) for 2 h at 60 W, the gel exposed on a Storage Phosphor Screen and imaged on a Typhoon PhosphoImager (GE Healthcare). The RNA and T DNA were quantified using Image J software (NIH). Released RNA and DNA were compared to the total samples to calculate fractions remaining in the TTC from triplicate reactions. These fractions vs. time were fit to a first-order dissociation rate equation with a small fraction that remained bound to the beads (recalcitrant to release) (Fig. 3H). The recalcitrant fractions (both RNA and DNA) were 0.05 for the first replicate and 0.13 for the second and third replicates. One timepoint (30 s) was lost for replicate 1.

### *In vitro* transcription assay

DNA templates used for in vitro transcription assays contains the T5-N25 promoter, a coding region, and the λ_tR2_ terminator. The DNA template was prepared by PCR primer extension using the single-stranded ntDNA (5’-TCATAAAAAATTTATTTGCTTTCAGGAAAATTTTTCTGTATAATAGATTCATAAATTTGAGAGAGGAGTTTAAATCCAGGCCTGCTGGTAA CGCAGGCCTTTTTATTTGGATCCCCGGGTAGAATTCG-3’; 1 μM; RuiMian) as template and tDNA (5’-CGAATTCTACCCGGGGATCCAAATAAAAAGGCCTGCGATT ACCAGCAGGCCTGGATTTA TGATCCCCGAGGAGAAGCAGAGGTACC-3’; 2 μM; Sangon Biotech) as the primer in a thermal cycler. The efficiency of primer extension was confirmed on a 1.5% agarose gel and the extended dsDNA were further purified by a Gel Extraction Kit (Omega Bio-Tek). The pre-melted DNA templates were prepared by the same procedure using tDNA primer with non-complementary sequences at the specified positions (Figs 3G, S8A, and S8B).

Reaction mixture (20 µL) in transcription buffer (40 mM Tris-HCl, pH 8.0, 75 mM NaCl, 5 mM MgCl_2_, 12.5% Glycerol, 2.5 mM DTT, and 50 μg/ml BSA) containing RNAP holo-enzyme (50 nM) and DNA template (50 nM) were incubated for 10 min at 37 °C. RNA synthesis was initiated by addition of 1.2 µL NTP mixture (5 µM ATP, 5 µM GTP, 0.05 µM UTP and 0.55 µM [α-^32^P]UTP (45 Bq/fmol), final concentration) for 10 min at 37 °C to obtain TECs halted at U25. Subsequently, 1µL Heparin (50 µg/mL; final concentration) was added to only allows single-round transcription. RNA extension was resumed by addition of 1 µL NTP mixture (5 µM ATP, 5 µM CTP, 5 µM GTP, and 5µM UTP; final concentration). Reactions were terminated by adding 5 μL loading buffer, boiled at 95 °C for 5 min, and cooled down in ice for 5 min. The RNA transcripts were separated by 12% urea-polyacrylamide slab gels (19:1 acrylamide/bisacrylamide) in 90 mM Tris-borate, pH 8.0 and 0.2 mM EDTA and analyzed by storage-phosphor scanning (Typhoon; GE Healthcare, Inc.). Promoter-based template DNA sequences with a λP_R_ promoter, C-less cassette, and wild-type or variant λ*t*_R2_ or ft_500_ terminators for experiments shown in Figs. S3 and S8C were PCR-amplified using primers flanking both ends and gel-purified using Qiagen Qiaquick purification reagents. NusA protein was purified as described previously^50^. Core RNAP was incubated with σ^70^ for 30 min at 37 °C to form holo-RNAP. To initiate transcription, holo-RNAP (31.25 nM) was incubated with template DNA (25 nM), ApU (150 µM), ATP + UTP (both at 2.5 µM), and 2 µM [α-^32^P]GTP (54.5 Ci/mmol) in EC buffer (10 mM Hepes, pH 8.0, 50 mM KGlutamate, 10 mM MgOAc, 0.1 mM EDTA, 5 µg acetylated BSA/mL and 1 mM DTT) for 5 min at 37 °C to form a halted complex at A26. Transcription was then restarted by adding a mastermix containing NTP mix (A+C+G+U), Heparin and KGlutamate at a final concentration of 150 µM, 50 µg/mL and 100 mM respectively. Reactions were stopped and products were separated as described above. The termination efficiencies were calculated from three independent replicates.

For experiments shown in Fig. S3F, ECs were reconstituted 15 nucleotides upstream from the U8 termination site by incubation of 25 nM RNA23, 50 nM T DNA, and 50 nM *E. coli* core RNAP in 50 µL EC buffer (10 mM Hepes, pH 8.0, 50 mM KGlutamate, 10 mM MgOAc, 0.1 mM EDTA, 5 µg/mL Acetylated BSA and 1 mM DTT) for 5 min at 37 °C. Non-template DNA (125 nM) was added and the mixture was incubated for an additional 5 min at 37 °C. Heparin (50 µg/mL) was added to the mixture to prevent core RNAP from rebinding to nucleic acids. The RNA was extended to A25 by a reaction with ATP + CTP (2.5 µM final for both) followed by radiolabeling to G27 by reacting with 0.037 µM [α-^32^P]GTP (3000 Ci/mmol). The RNA was then extended to the termination site by addition of 150 µM NTP mix.

## Data availability

The cryo-EM map and coordinates were deposited in Protein Data Bank and Electron Microscopy Data Bank (TTC-pause: 7YP9 and EMD-33996; TTC-hairpin: 7YPA, EMD-33997; TTC-release: 7YPB, EMD-33998).

## Acknowledgments

The Work was supported by the Basic Research Zone Program of Shanghai JCYJ-SHFY-2022-012 (YZ), National Key Research and Development Program of China 2018YFA0900701 (Y.Z.), Strategic Priority Research Program of the CAS XDB29020000 (Y.Z.), and US National Institute of General Medical Sciences, NIH (GM38660) (R.L.). We thank Dr. Liangliang Kong, Dr. Fangfang Wang, Dr. Guangyi Li, and Dr. Jialin Duan at the cryo-EM center of NFPS in Shanghai, Dr. Shenghai Chang at the cryo-EM center of Zhejiang University.

## Author contributions

L.Y. collected the cryo-EM data, solved the cryo-EM structures. L.Y., E.O., C.Y. and R.M. performed biochemical experiments. S.J., L.S., X.W., D.H., Y.Z. and Y.F. assisted in structure determination. R.L. and Y.Z. designed experiments, analyzed data, and wrote the manuscript.

## Materials & Correspondence

Correspondence and requests for materials should be addressed to Y.Z. or R.L.

